# Controlled Hydrogen Sulfide Delivery to Enhance Cell Survival in Bone Tissue Engineering

**DOI:** 10.1101/2022.11.14.516486

**Authors:** Soheila Ali Akbari Ghavimi, Trent J. Faulkner, Rama Rao Tata, Ethan S. Lungren, Rui Zhang, Erin E. Bumann, Bret D. Ulery

## Abstract

The increased local concentration of calcium ions (Ca^2+^) and phosphate (P_i_), a natural body process for bone healing and remodeling, as well as local delivery of these ions as signaling molecules by synthetic bone graft substitutes, may lead to cytotoxic ion levels that can result in Ca^2+^/ P_i_ mitochondria overload, oxidative stress, and cell death. In this research, the effect of H_2_S as a cytoprotective signaling molecule to increase the tolerance of mesenchymal stem cells (MSCs) in the presence of cytotoxic level of Ca^2+^/P_i_ was evaluated. Different concentrations of sodium hydrogen sulfide (NaSH), a fast-releasing H_2_S donor, were exposed to cells in order to evaluate the influence of H_2_S on MSC proliferation. The results suggested that a range of NaSH (*i.e*., 0.25 - 4 mM NaSH) was non-cytotoxic and could improve cell proliferation and differentiation in the presence of cytotoxic levels of Ca^2+^ (32 mM) and/or P_i_ (16 mM). To controllably deliver H_2_S over time, a novel donor molecule in thioglutamic acid (GluSH) was synthesized and evaluated for its H_2_S release profile. Excitingly, GluSH successfully maintained cytoprotective level of H_2_S over 7 days. Furthermore, MSCs exposed to cytotoxic Ca^2+^/P_i_ concentrations in the presence of GluSH were able to thrive and differentiate into osteoblasts. These findings suggest that the incorporation of a sustained H_2_S donor such as GluSH into CaP-based bone substitutes can facilitate considerable cytoprotection making it an attractive option for complex bone regenerative engineering applications.

## INTRODUCTION

Bone defects are the second leading cause of disability affecting more than 1.7 billion people worldwide.^1^ When bone fractures occur, free radicals and reactive oxygen species (ROS) are generated within the damaged tissue causing an imbalance between these and antioxidants damaging cellular macromolecules and altering their functions through oxidative stress.^2^ This oxidative stress causes Ca^2+^/P_i_ influx into the cytoplasm from the extracellular environment followed by Ca^2+^/P_i_ passage into the cell mitochondria. The trauma associated with fractures or other major bone injuries also damages the blood supply, resulting in local hypoxia.^3^ An increase in intracellular Ca^2+^ levels is a primary response of many cell types to hypoxia.^4^ Finally, after considerable bone tissue damage, Ca^2+^/P_i_ concentrations in the blood and urine will decrease^5^ while levels local to the defect site will increase causing the formation of a soft callus which is necessary for bone remodeling.^6^ These increases in extracellular Ca^2+^/P_i_ concentrations can cause an intracellular overload.^7^ All three of these phenomena contribute to Ca^2+^/P_i_ mitochondrial overload disrupting and accelerating cellular metabolism leading to cell death.^3^ In addition, mitochondrial Ca^2+^/P_i_ overload has been found to lead to greater ROS production^4^ which further increases oxidative stress leading to even greater intracellular Ca^2+^/P_i_ concentrations.

While bones are capable of remodeling themselves, reconstruction of critical-sized bone defects is challenging as intervention is required to facilitate adequate healing.^8^ Since the body’s natural reaction to bone tissue damage is to locally increase Ca^2+^/P_i_ concentrations, many regenerative engineering approaches leverage these ions as signaling molecules in their design. Bioactive ceramics, especially calcium phosphates (CaPs), have been widely used in bone regeneration because of their similarity to native bone mineral content.^9, 10^ Once immersed in an aqueous solution, CaPs undergo dissolution and precipitation as the result of ion transfer at the solid–liquid interface yielding a net release of calcium ions (Ca^2+^) and phosphate ions (H_2_PO_4_^-^, HPO_3_^2-^, or PO_4_^3-^ - P_i_) from the material.^11^ These ions have been found to facilitate bone regeneration.^12, 13^ Ca^2+^ and P_i_ osteoinductivity exists in a concentration range termed the therapeutic window where enough ions are present to facilitate stem cell osteogenic differentiation, but not too many to overwhelm the cells inducing their death. Our previous efforts have shown that Ca^2+^ concentrations of 32 mM or more and P_i_ concentrations of 16 mM or more are cytotoxic for mesenchymal stem cells *in vitro*^14^ and are the limits that can be used for biomaterials-based bone regeneration engineering applications^15,16^.

Improving cell tolerance to increased Ca^2+^/P_i_ concentrations can led to a broader and enhanced therapeutic window by limiting the cytotoxic effects of high concentration Ca^2+^/P_i_. One promising option to mitigate ion-mediated toxicity is hydrogen sulfide (H_2_S), a gasotransmitter signaling molecule^17^ that has been found capable of suppressing oxidative stress in mitochondria^18–20^ as well as moderating cellular oxygen consumption especially under hypoxic conditions.^21^ Unfortunately, there are only a few H_2_S donors commercially available with most research efforts utilizing sodium hydrosulfide (NaHS) or sodium sulfide (Na_2_S).^22^ As both of these rapidly dissociate in water, they yield an uncontrollable burst release of H_2_S that can be toxic to cells. L-cysteine is another option where H_2_S is enzymatically liberated from the donor molecule. This process is regrettably not externally controllable as enzymatic downregulation is tied to increased local H_2_S levels.^23^ Some synthetic H_2_S donors have been developed including GYY4137 for more sustained and hydrolytic cleavage, but the toxicity and clearance of the remaining small molecule backbone after H_2_S release has yet to be studied.^24, 25^ Zhou and colleagues developed H_2_S donors through thioacid substitution in amino acids (*i.e*., glycine and valine) to improve the compatibility, predictability, and degradation kinetics of a H_2_S donor as a cardioprotective reagent.^24^ While promising, the rapid release rate and lack of a conjugatable chemical group makes thioglycine and thiovaline difficult to incorporate into various biomaterials for sustained, localized H_2_S release to supplement site-directed Ca^2+^/P_i_ osteoinductivity. To address this issue, we have designed and synthesized a novel H_2_S donor with a conjugatable carboxylic acid (*i.e*., thioglutamic acid - GluSH) and evaluated its cytoprotective effect on mesenchymal stem cells exposed to cytotoxic Ca^2+^/P_i_ concentrations *in vitro*.

## EXPERIMENTAL

### Preparation of H_2_S, Ca^2+^, and P_i_ Containing Media

H_2_S stock solution (pH = 7.4) was prepared by dissolving 512 mM of NaSH (Sigma-Aldrich) in distilled, deionized water (ddH_2_O) at 37 °C. Ca^2+^ stock solution (pH = 7.4) was prepared by dissolving 512 mM calcium chloride (CaCl_2_; Sigma-Aldrich), 25 mM 4-(2-Hydroxyethyl)piperazine-1-ethanesulfonic acid (HEPES; Sigma-Aldrich), and 140 mM sodium chloride (NaCl; Sigma-Aldrich) in ddH_2_O at 37 °C. P_i_ stock solution (pH = 7.4) was prepared by dissolving disodium hydrogen phosphate dihydrate (Na_2_HPO_4_ • 2 H_2_O; Sigma-Aldrich) and sodium dihydrogen phosphate dihydrate (NaH_2_PO_4_ • 2 H_2_O; Sigma-Aldrich) at a ratio of 4:1 Na_2_HPO_4_/NaH_2_PO_4_, 25 mM HEPES, and 140 mM NaCl in ddH_2_O at 37 °C. These stock solutions were made fresh for each experiment and sterilized by 0.22 μm syringe filter. Individual solution concentrations of 0.25, 0.5, 1, 2, 4, 8, 16, 32, and 64 mM H_2_S, 32 mM Ca^2+^, and 16 mM P_i_, as well as combined solution concentrations of 32 mM: 1 mM Ca^2+^:H_2_S, 16 mM: 1 mM P_i_:H_2_S, and 32 mM: 16 mM: 1 mM H_2_S:Ca^2+^:P_i_:H_2_S were prepared by diluting stock solutions in growth media consisting of Dulbecco’s modified Eagle’s medium (DMEM, Invitrogen) supplemented with 10% fetal bovine serum (FBS, Invitrogen) and 1% Penicillin-streptomycin (Pen-Strep, Invitrogen).

### Cell Culture and Seeding

Murine mesenchymal stem cells (MSCs) were purchased from Cyagen and initially cultured in T-75 cell culture flasks (Corning) in growth medium at 37 °C in a humidified incubator supplemented with 5% CO_2_. Media was changed every 48 h until cells approached ~ 80% confluency after which they were dissociated using a 0.05% trypsin-EDTA (Invitrogen) solution. Detached MSCs were counted by hemocytometer and passed to new T-75 flasks at a splitting ratio of 1:4 or 1:5 dependent on cell count. After the 5th passage, cells were used for *in vitro* bioactivity studies. Tissue cultured polystyrene 24-well plates (Corning) were seeded with 30,000 cells/well and exposed to growth media alone as a negative control or growth media containing 0.25, 0.5, 1, 2, 4, 8, 16, 32, 64, 128, or 256 mM NaSH, or 32 mM Ca^2+^ (Ca_32_), or 16 mM P_i_ (P_16_). Additional studies utilized growth media containing 32 mM: 16 mM Ca^2+^:P_i_ (Ca_32_/P_16_), 32 mM glutamic acid (Glu_32_), 32 mM GluSH (GluSH_32_), 32 mM: 1 mM Ca^2+^:NaSH (Ca_32_/NaSH_1_), 16 mM: 1 mM P_i_:NaSH (P_16_/NaSH_1_), 32 mM: 16 mM: 1 mM Ca^2+^:P_i_:NaSH (Ca_32_/P_16_/NaSH_1_), 32 mM: 32 mM Ca^2+^:Glu (Ca_32_/Glu_32_), 16 mM: 32 mM P_i_:Glu (P_16_/Glu_32_), or 16 mM: 32 mM: 32 mM Ca^2+^:P_i_:Glu (Ca_32_/P_16_/Glu_32_), 32 mM: 32 mM Ca^2+^:GluSH (Ca_32_/GluSH_32_), 16 mM: 32 mM P_i_:GluSH (P_16_/GluSH_32_), or 32 mM: 16 mM: 32 mM Ca^2+^:P_i_:GluSH (Ca_32_/P_16_/GluSH_32_) loaded into a Transwell membrane insert (Corning). MSCs were cultured with these solutions for up to 7 days at 37 °C in a humidified incubator supplemented with 5% CO_2_ and the media containing various compounds were added fresh every 2 days. After day 7, cells exposed to high Ca^2+^ and P_i_ concentrations were treated with 16 mM Ca^2+^ (Ca_16_), 8 mM P_i_ (P_8_), or 16 mM: 8 mM Ca^2+^:P_i_ (Ca_16_/P_8_) based on their group without any additional H_2_S releasing molecules with the media containing appropriate ion concentrations changed every 2 days. Cell proliferation, viability, alkaline phosphatase (ALP) activity, and mineralization were assessed at 1, 3, 7, and 14 days.

### Proliferation Assay

Cell proliferation was determined using the Quanti-iT PicoGreen dsDNA Assay (Thermo Fisher Scientific). At each endpoint, the samples were rinsed with phosphate buffered saline (PBS) and exposed to 1% Triton X-100 (Sigma-Aldrich) followed by three freeze-thaw cycles in order to lyse the cells. Lysates were diluted with TE buffer (200 mM Tris-HCL, 20 mM EDTA, pH 7.5) and mixed with PicoGreen reagent according to the manufacturer’s protocol. A BioTek Cytation 5 fluorospectrometer plate reader was utilized to measure the fluorescence of each sample (*ex*. 480 nm, *em*. 520 nm) and the cell number was calculated using a MSC standard curve (0 - 200,000 cells/mL).

### Viability Assay

Cell viability was evaluated at each time point using an MTS Cell Proliferation Colorimetric Assay Kit (BioVision). MTS reagent (20 μL) was added to growth media (500 μL) followed by 4 h incubation at 37 °C in a humidified incubator supplemented with 5% CO_2_. Absorbance of each sample was measured at 490 nm using a plate reader. Cell viability was reported as the ratio of absorbance in the experimental groups compared to the growth media negative control.

### Alkaline Phosphatase Activity Assay

Cell ALP activity was quantified at each time point using an Alkaline Phosphatase Assay Kit (BioVision). In brief, 20 μL of cell lysate was combined with 50 μL of p-nitrophenyl phosphate (pNPP) solution in assay buffer. The mixture was incubated for 1 h at room temperature away from light. The reaction was stopped by adding 20 μL of the stop solution and the absorbance of the solution was measured at 405 nm using a plate reader. To eliminate any background effects, 1% Triton X-100 was incubated with pNPP, exposed to stop solution in assay buffer after 1 h, and its absorbance deducted from sample absorbance. The absorbance was converted to content of dephosphorylated p-nitrophenyl (pNP) using a pNP standard curve (0 - 20 nmol/mL) which was dephosphorylated using excess ALP Enzyme. ALP activity was reported as the pNP content normalized by cell count.

### Mineralization Assay

Cell-based mineral deposition was measured using an Alizarin red assay. At each time point, the media was removed after which the cells were washed with ddH_2_O and fixed in 70% ethanol for 24 h. The ethanol was removed and the samples were incubated in 1 mL of 40 mM Alizarin Red solution (Sigma-Aldrich) for 10 minutes. The samples were rinsed with ddH_2_O several times to make sure all non-absorbed stain was removed. Absorbed Alizarin Red was desorbed using 1 mL of a 10% cetylpyridinium chloride (CPC, Sigma-Aldrich) solution after which stain concentration was measured at 550 nm using a plate reader. Absorbance of each sample was converted to the concentration of absorbed Alizarin Red using a standard curve (0 - 0.2740 mg/mL). Samples above the standard curve linear range were diluted with CPC solution until a reading in the linear range was obtained. Cell-based mineral deposition was calculated by subtracting mineralization found in blank wells exposed to the same experimental conditions. All results were normalized by cell count.

### GluSH Synthesis

The three-step synthesis process is detailed in **Scheme 1**.

**Scheme 1.**
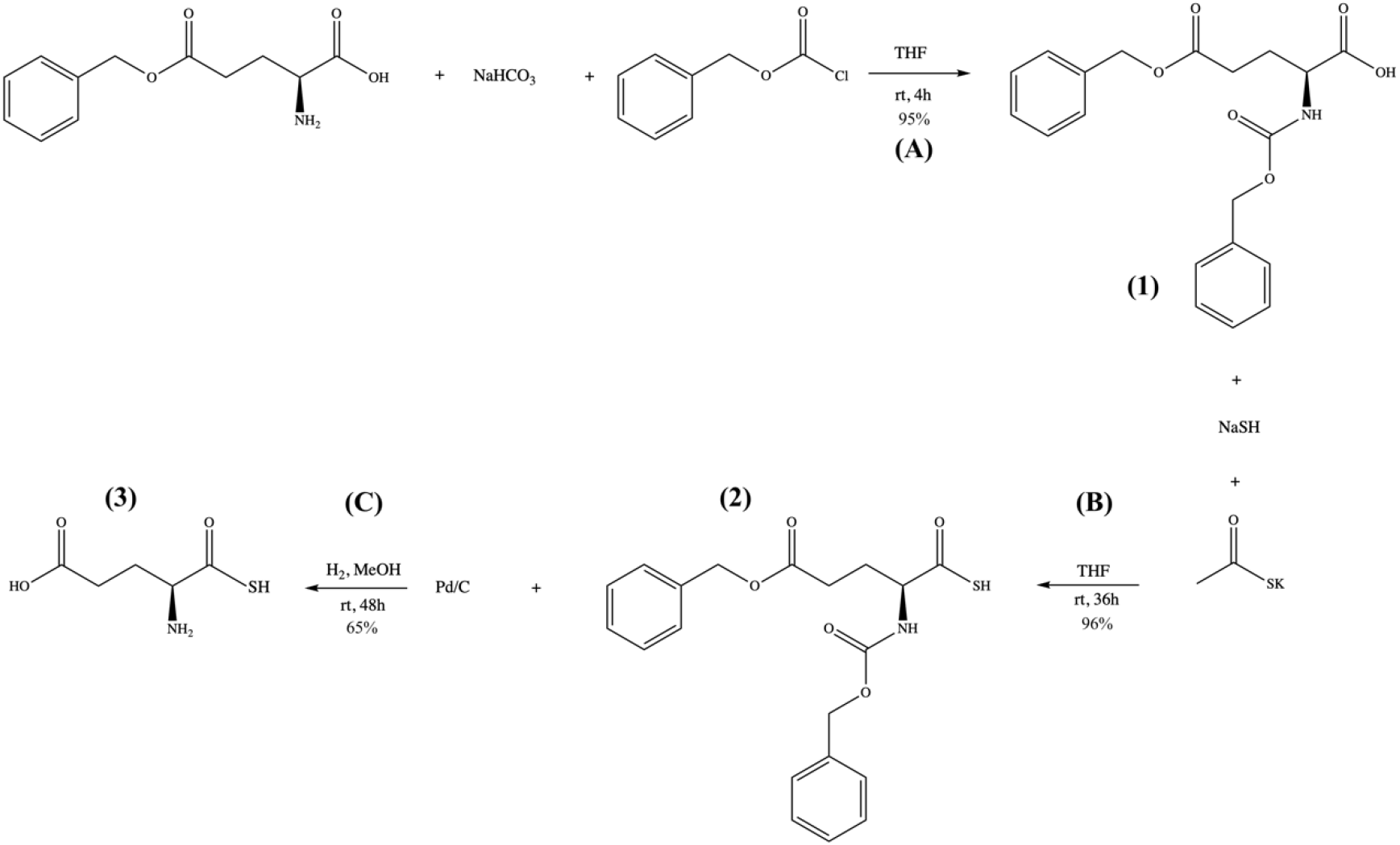
GluSH synthesis process.

#### Reaction A

Sodium bicarbonate (NaHCO_3_; Sigma-Aldrich) (1.5 mmol) was dissolved in water (50 mL) at room temperature after which 10 mL of dry tetrahydrofuran (THF; Sigma-Aldrich) was added. L-glutamic acid 5-benzyl ester (Alpha Aesar) (4.2 mmol) was dissolved in the reaction mixture followed by dropwise addition of benzyl chloroformate (Sigma Aldrich) (6.5 mL). After 4 hours stirring under an argon atmosphere, the reaction mixture turned to a clear solution. To remove the unreacted benzyl chloroformate, the reaction mixture was washed with diethyl ether (60 mL x 2). The aqueous layer was then acidified (pH ~ 2) with 1 mM hydrochloric acid (HCl). The mixture was then extracted with ethyl acetate (60 mL x 2), dried over anhydrous sodium sulfate (Na_2_SO_4_), and evaporated under reduced pressure using a rotary evaporator (Buchi). The final product, N-benzyloxycarbonyl-L-glutamic acid 5-benzyl ester **(1)**, was a white powder and used without further purification. This product was dissolved in deuterated chloroform for ^1^H-NMR spectroscopy analysis (**Fig. S1**): ^1^H NMR (500 MHz, CDCl_3_) δ 7.35-7.27 (m, 10H), 5.56-5.54 (m, 1H), 5.11 (m, 4H), 4.43-4.40 (m, 1H), 2.51-2.45 (m, 2H), 2.27-2.24 (m, 1H), 2.04 (m, 1H).

#### Reaction B

N-benzyloxycarbonyl-L-glutamic acid 5-benzyl ester (**(1)**, 10 mmol) were dissolved in 15 mL of dry THF in a round bottom flask. The solution was then treated with NaSH (20 mmol) and thioacetic acid (10 mmol) at room temperature. After the reaction mixture was stirred in open air for 48 hours, the solvent was evaporated under reduced pressure using a rotary evaporator. The residue was diluted with water (50 mL) and acidified with 1 mM HCl (pH ~ 2). This aqueous solution was extracted with ethyl acetate (30 mL x 3). The combined organic layer was washed with brine, dried over anhydrous Na_2_SO_4_, and evaporated under reduced pressure using a rotary evaporator. The final product was purified using a silica gel column chromatography and the pure N-benzyloxycarbonyl-L-thioglutamic acid 5-benzyl ester **(2)** was eluted at 40% ethyl acetate in hexane as a yellow oil. This product was dissolved in deuterated chloroform for ^1^H-NMR spectroscopy analysis (**Fig. S2**): ^1^H NMR (500 MHz, CDCl_3_) δ 7.35-7.27 (m, 10H), 5.53-5.51 (m, 1H), 5.12 (m, 4H), 4.46-4.45 (m, 1H), 2.56-2.48 (m, 2H), 2.31-2.29 (m, 1H), 2.07 (m, 1H).

#### Reaction C

N-benzyloxycarbonyl-L-thioglutamic acid 5-benzyl ester (**(2)**, 10 mmol) was dissolved in methanol (40 mL) after which 10 wt % palladium on carbon (Pd/C; Sigma-Aldrich) was added to the mixture as a catalyst for the hydrogenation reaction. The reaction flask was sealed and vacuum purged prior to the addition of hydrogen gas. The reaction was stirred for 48 hours under hydrogen at room temperature. The reaction mixture was then dissolved in water and vacuum filtered. The clear filtrate was then evaporated under reduced pressure using a rotary evaporator which left a yellow powder of L-thioglutamic acid (GluSH, **(3)**). This product was dissolved in deuterium oxide for ^1^H-NMR spectroscopy analysis (**Fig. S3**): ^1^H NMR (500 MHz, D2O) δ 3.96-3.79 (m, 1H), 2.57-2.56 (m, 2H), 2.16 (m, 2H).

### H_2_S Release Study

H_2_S release from GluSH was measured in ddH_2_O at 37 °C over 7 days. GluSH loaded into a Transwell membrane insert (Corning) was placed in centrifuge tubes containing Hank’s buffer saline solution (HBSS) supplemented with excess amounts of bicarbonate (1.5 mol per 1 mol GluSH). Each centrifuge tube was sealed with vacuum glue and parafilm. At certain time points, 1 mL of the release solution were extracted from the reaction tube using a needle and replaced with 1 mL of HBSS and bicarbonate solution. The solutions then immediately assayed to determine their H_2_S concentration using a fluorescent method measuring the formation of thiobimane.^26^ Briefly, dibromobimane (Sigma-Aldrich) was dissolved in HBSS at a concentration of 500 μM and incubated with the test samples at room temperature for 5 minutes after which florescence (*ex*. 340 nm, *em*. 465 nm) was measured. A standard curve was created by dissolving different concentrations of NaSH in the 500 μM dibromobimane in HBSS solution, incubating at room temperature for 5 minutes, and measuring the florescence. The concentration of the H_2_S in each time point were then calculated by comparing the test sample values to the NaSH standard curve.

### Statistical Analysis

JMP software was used to make comparisons between groups with Tukey’s HSD test specifically utilized to determine pairwise statistical differences (p < 0.05). The statistical analysis results are reported in the supporting information section. Groups that possess different letters have statistically significant differences in mean whereas those that possess the same letter have means that are statistically insignificant in their differences.

## RESULTS AND DISCUSSION

### H_2_S Cytotoxicity

Proliferation and viability of MSCs exposed to different concentrations of NaSH are shown in **Fig. 1**. MSCs seeded on tissue cultured plastic (Ctrl) and incubated in growth media increased to 7 times initial cell seeding number over 14 days and cells exposed to concentrations of 4 mM NaSH or less expanded to more than 8 times initial cell seeding number (**Fig. 1A**). The cell number for those exposed to 0.5 and 1 mM of NaSH were statistically significantly higher than control group at day 14 (**Table S1**). On the other hand, proliferation of MSCs exposed to 8 mM NaSH or more were statistically significantly lower than those provided growth medium supplemented with 0 - 4 mM NaSH (**Table S1**). The mildly mitogenic behavior of H_2_S at lower concentrations is likely due to its ability to increase intracellular levels of cyclic guanosine monophosphate (cGMP),^27^ which is known to stimulate stem cell proliferation.^28^ H_2_S can also inhibit cytochrome c oxidase activity and cause oxidative stress^29^ or even directly cause a radical-associated DNA damages,^30^ which is likely responsible for the cell death found at higher H_2_S concentrations.

**Fig. 1.**
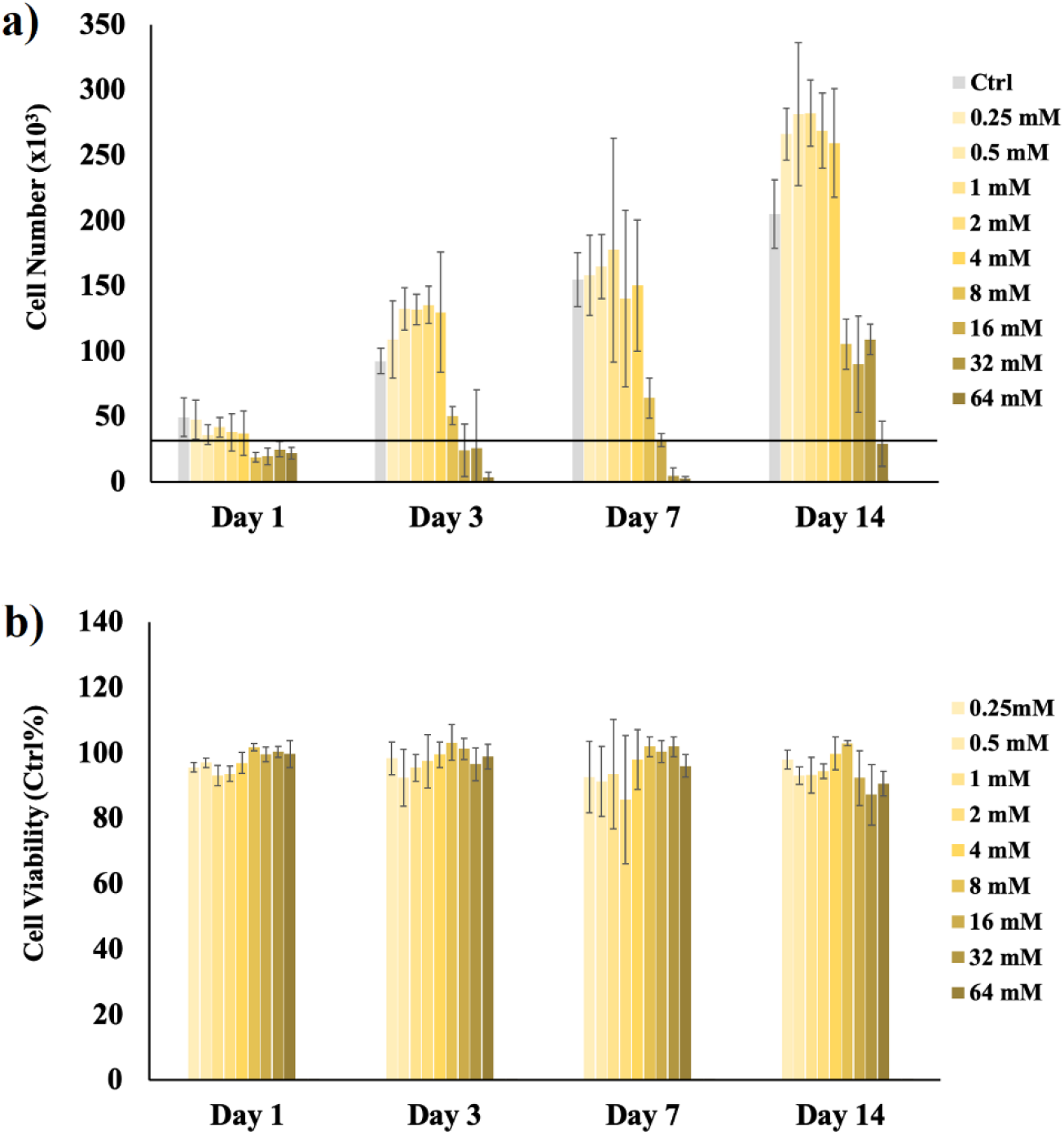
Proliferation and viability of MSCs exposed to different concentrations of NaSH supplemented media. **(a)** Cell count was measured by the Quanti-iT PicoGreen Assay over 14 days of cell culture in media supplemented with 0 - 256 mM NaSH for 7 days and then culture media with 0 mM NaSH for the next 7 days. The black line indicates the original cell seeding number (*i.e*., 30,000). **(b)** An indirect measure of cell metabolism and viability was determined by NAD(P)H activity over 14 days using a MTS assay for MSCs exposed to 0 - 256 mM NaSH for 7 days and then culture media with 0 mM NaSH for the next 7 days. Statistical analysis of the data is available in the supplementary information (**Table S1** and **Table S2**).

The results of the MTS assay show that MSCs exposed to 4 mM NaSH or less were over 90% as viable as the control group through 14 days. Interestingly, MSCs subj ected to 8 mM NaSH or more were still more than 80% viable as the control group through 14 days (**Fig. 1B**). Taken with the proliferation results, these data indicate that though high concentrations of NaSH adversely affect MSC proliferation, the surviving cells are highly metabolically active over the 14 days of the study.

### Cytoprotective Effect of H_2_S at High Ca^2+^ and/or P_i_ Concentrations

The cytotoxicity evaluation of NaSH revealed that high NaSH concentration (*i.e*., 8 mM or more) can decrease cell proliferation. Since there was no statistically significantly difference between cell proliferation and viability of the MSCs exposed to 0.25 to 4 mM NaSH, the rest of this research explored the cytoprotective effect of 1 mM NaSH (NaSH_1_) when co-delivered with higher concentrations of Ca^2+^ and/or P_i_. Our previous research indicates that media supplemented with more than 16 mM Ca^2+^ and/or 8 mM P_i_ can be cytotoxic adversely affecting MSC proliferation and viability.^14^ Therefore, 32 mM Ca^2+^ (C32 - cytotoxic concentration), 16 mM P_i_ (P_16_ - cytotoxic concentration), 1 mM NaSH (NaSH_1_ - non-cytotoxic concentration), and combinations of Ca^2+^, P_i_, and NaSH_1_ were used to explore the cytoprotective effect of hydrogen sulfide in the initial presence of excess calcium and phosphate ions. After 7 days of high concentration exposure, MSCs were exposed to inductive and non-cytotoxic Ca^2+^ (*i.e*., Ca_16_) and P_i_ (*i.e*., P_8_) concentrations without NaSH from day 7 to day 14 to better mimic the conditions within the bone fracture site.

Proliferation and viability of cells exposed to different combinations of ionic and gasotransmitter signaling molecules is demonstrated in **Fig. 2**. MSCs cultured with Ca_32_ and P_16_ showed less proliferation over 14 days as compared to those grown under control conditions, which is due to the likely cytotoxic effects and the possible osteoinductivity of high Ca^2+^ and P_i_ concentrations (**Fig. 2A**). However, when the MSCs were also supplied with NaSH_1_, their proliferation was statistically significantly greater through 14 days (**Table S3**). This result demonstrate the positive effect of NaSH on the proliferation of cells exposed to cytotoxic level of Ca^2+^ and P_i_. It is known that increases in intracellular Ca^2+^ promote mitochondrial calcium uptake resulting in the loss of mitochondrial membrane potential which is interpreted as a danger signal initiating apoptosis.^31^

**Fig. 2.**
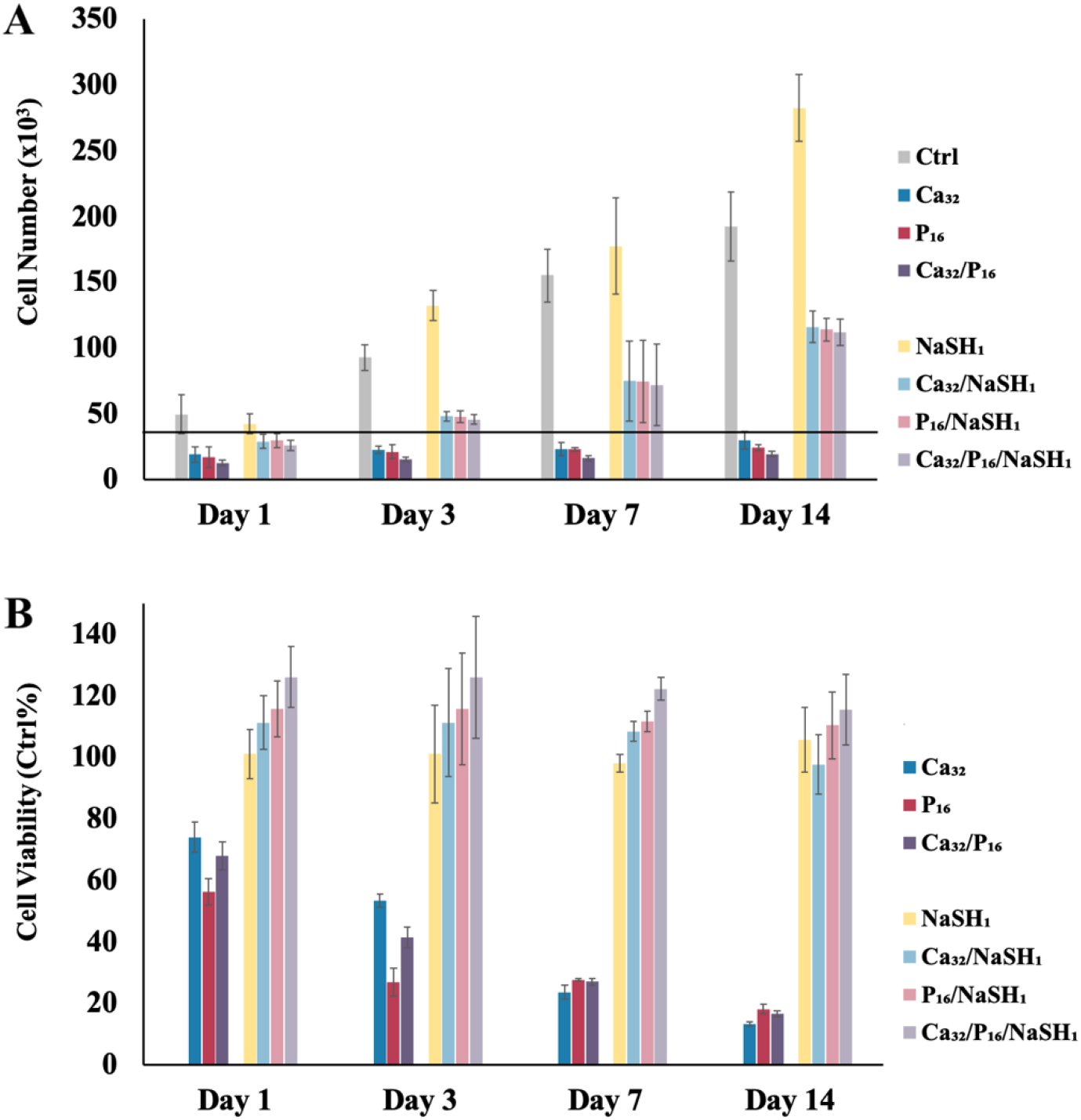
Proliferation and viability of MSCs exposed to different combinations of Ca^2+^, P_i_, and/or NaSH supplemented media. **(a)** Cell count was measured by the Quanti-iT PicoGreen Assay over 14 days of cell culture in media supplemented with no signaling molecules (Ctrl), 32 mM Ca^2+^ (Ca_32_), 16 mM P_i_ (P_16_), Ca_32_/P_16_, 1 mM NaSH (NaSH_1_), Ca_32_/NaSH_1_, P_16_/NaSH_1_, or Ca_32_/P_16_/NaSH_1_ for the first 7 days. This was followed by the cells being exposed for the next 7 days to no signaling molecules or half the Ca^2+^ and/or P_i_ concentrations (*i.e*., Ca_16_ and/or P_8_) they were originally cultured in. The black line indicates the original cell seeding number (*i.e*., 30,000). **(b)** An indirect measure of cell metabolism and viability was determined by NAD(P)H activity over 14 days using an MTS assay. MSCs were exposed to Ca_32_, P_16_, and/or NaSH_1_ and combination of these molecules (Ca_32_/P_16_, Ca_32_/NaSH_1_, P_16_/NaSH_1_, and Ca_32_/P_16_/NaSH_1_) for the first 7 days followed by the aforementioned decrease in Ca^2+^ and/or P_i_ concentrations for the respective groups. All data were normalized against the results determined for MSCs given non-supplemented media. Statistical analysis of the data is available in the supplementary information (**Table S3** and **Table S4**).

Addition of NaSH_1_ as a H_2_S donor can help regulate cytosolic Ca^2+^ levels and mitochondrial calcium uptake,^32^ resulting in higher cell survival compared to MSCs exposed to a high Ca^2+^ concentration alone. Higher P_i_ concentrations can dysregulate the mitochondrial permeability transition pore (MPTP) initiating a pro-apoptotic cascade^33^ or further enhancing ongoing apoptosis.^34^ By reducing oxidative stress, H_2_S indirectly protects the mitochondria from becoming damaged and activating pro-apoptotic signaling pathways.^35^

Cell viability results reveal that the metabolic activity of MSCs exposed to the Ca_32_, P_16_, and Ca_32_/P_16_ were statistically significantly lower than Ca_32_/NaSH_1_ and P_16_/NaSH_1_, and Ca_32_/P_16_/NaSH_1_ through 14 days (**Fig. 2B** and **Table S3**). High Ca^2+^ and/or P_i_ concentrations can even lead to viability less than 20% when compared to the control group whereas supplementing these same groups with NaSH_1_ preserved cell viability to over 90% of control. To investigate viability, the MTS assay was used which measures cell viability by indirectly determining NAD(P)H-dependent mitochondrial dehydrogenase activity essential to cell metabolism and proliferation oxidation/reduction reactions.^36^ The previously mentioned mitochondrial dysregulation due to Ca_32_ and P_16_ that limited cell proliferation understandably negatively impacted cell dehydrogenase activity as well.

ALP activity and cell-based mineralization of MSCs cultured with Ca_32_ or P_16_ with or without NaSH_1_ is described in **Fig. 3**. MSC osteogenic differentiation is known to proceed through 3 stages: proliferation, maturation, and mineralization. ALP is an enzyme that is expressed during the beginning of the osteogenic maturation period.^37^ The ALP results presented were normalized by cell number to focus on cell differentiation independent of proliferation. MSCs exposed to media (Ctrl) and media supplemented with NaSH_1_ showed background levels of ALP expression while cells subjected to Ca_32_, P_16_, Ca_32_/P_16_, Ca_32_/NaSH_1_, P_16_/NaSH_1_, and Ca_32_/P_16_ /NasSH1 showed statistically significant increased levels of ALP expression compared to control (**Fig. 3A**, **Table S3**). Also, cells cultured with Ca_32_ and P_16_ supplemented with NaSH_1_ (*i.e*., Ca_32_/NaSH_1_, P_16_/NaSH_1_, and Ca_32_/P_16_ /NasSH1) showed statistically significantly higher levels of ALP expression compared to those without NaSH_1_ (*i.e*., Ca_32_, P_16_, and Ca_32_/P_16_) at each time point assessed (**Fig. 3A**, **Table S3** and **S4**). The low ALP activity in MSCs exposed to Ca_32_ and/or P_16_ alone is likely due to their diminished viability which when modulated by NaSH_1_ can improve overall ALP activity considerably. It is recently been reported that H_2_S can also promote osteoblast differentiation at sites of bone regeneration by triggering deposition of inorganic mineral matrix and promoting expression of osteogenic genes in human MSCs.^38^ However, our results did not show improved ALP activity after exposure of MSCs to NaSH_1_ alone compared to control group over the course of 14 days suggesting H_2_S can assist other inductive molecules (*i.e*., Ca^2+^ and P_i_) though may not be osteoinductive in its own right.

**Fig. 3.**
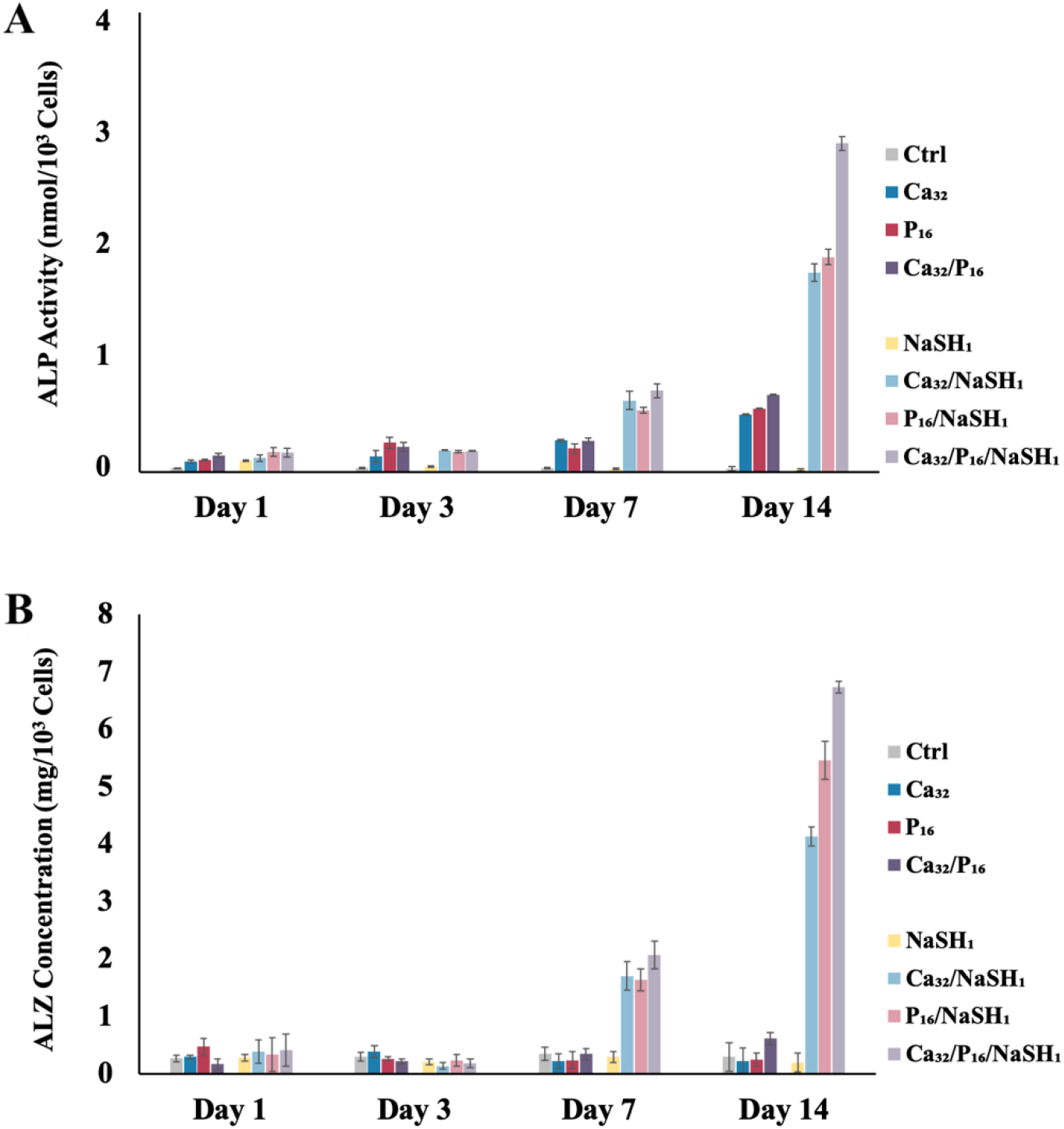
ALP activity and cell-based mineralization of MSCs exposed to different combinations of Ca^2+^, P_i_, and/or NaSH supplemented media. **(a)** Maturation enzyme activity was analyzed by an ALP Assay Kit over 14 days of cell culture in media supplemented with no signaling molecules (Ctrl), 32 mM Ca^2+^ (Ca_32_), 16 mM P_i_ (P_16_), Ca_32_/P_16_, 1 mM NaSH (NaSH_1_), Ca_32_/NaSH_1_, P_16_/NaSH_1_, or Ca_32_/P_16_/NaSH_1_ for the first 7 days. This was followed by the cells being exposed for the next 7 days to no signaling molecules or half the Ca^2+^ and/or P_i_ concentrations (*i.e*., Ca_16_ and/or P_8_) they were originally cultured in. **(b)** Alizarin red (ALZ) staining was used as an indirect measure of mineralization over 14 days of cell culture. MSCs were exposed to no signaling molecules (Ctrl), Ca_32_, P_16_, and/or NaSH_1_, and combination of these molecules (Ca_32_/P_16_, Ca_32_/NaSH_1_, P_16_/NaSH_1_, Ca_32_/P_16_/NaSH_1_) for the first 7 days followed by the aforementioned decrease in Ca^2+^ and/or P_i_ concentrations for the respective groups. Both ALP activity and ALZ mineralization data were normalized on a per cell basis. Statistical analysis of the data is available in the supplementary information (**Table S3** and **Table S4**).

In comparison the early-stage activity of ALP, matrix mineralization is a late-stage MSC osteogenic marker. In this research, acellular experiments were conducted in parallel using the same conditions allowing their results to be subtracted from the complementary cellular groups to eliminate solution-mediated mineralization. These cell-based mineralization results were then normalized by cell number to evaluate improved osteoinductivity by NaSH_1_ independent of its influence on proliferation. Evaluation of MSC mineralization revealed that Ca_32_/NaSH_1_, P_16_/NaSH_1_, and Ca_32_/P_16_/NaSH_1_ induced statistically significantly greater mineral deposition over time (**Table S4**) as well as greater than cells exposed to Ctrl and NaSH_1_ treatments (**Fig. 3B** and **Table S3**). Ca_32_, P_16_, and Ca_32_/P_16_ were found to limit mineralization instead of enhancing it over the Ctrl group over time (**Fig. 3B** and **Table S3** and **S4**) likely due to the toxicity of these treatments. The mineralization results were in agreement with the ALP activity data supporting the beneficial effects of H_2_S when supplementing high concentrations of osteoinductive ions.

### H_2_S Release from GluSH

In order to investigate the cytoprotective effect of sustained H_2_S delivery, GluSH was synthesized and tested for its ability to controllably release H_2_S over time. GluSH is a highly water soluble product that releases a sulfanyl ion (HS^-^) in the presence of bicarbonate by the chemical reaction outlined in **Fig. 4** which can then be protonated in water to yield H_2_S. HS^-^ is only produced from GluSH in the presence of bicarbonate, which is a molecule that can be readily found in the human body.^39^ Therefore, bicarbonate-supplemented pH and osmotic balancing HBSS was used to investigate H_2_S release from GluSH over time. GluSH quantity used in the experiment was varied to determine what concentration would allow the maximum H_2_S concentration release to not exceed the previously NaHS-determined cytotoxic level (*i.e*., 4 mM). **Fig. 5** summarizes the results of H_2_S production from 32 mM GluSH over 7 days. As shown in **Fig. 5A**, GluSH released around 50% of its total H_2_S generating payload within the first day. H_2_S continued to be generated through day 5 after which little more was alleviated from GluSH. When the data was converted to H_2_S concentration, GluSH was found to controllably release its payload within the non-cytotoxic range (*i.e*., 0.25 - 4 mM) for 7 days (**Fig. 5B**).

**Fig. 4.**
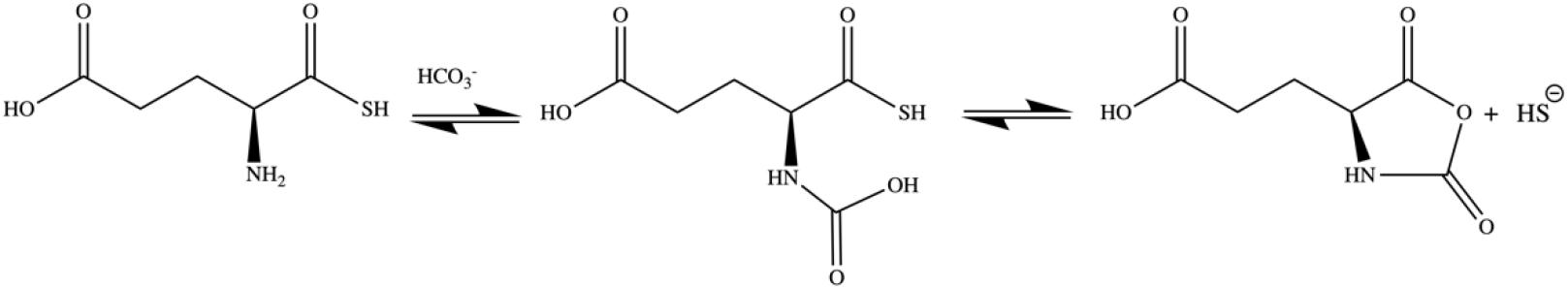
Sulfanyl ion (HS^-^) release mechanism from GluSH in the presence of bicarbonate. The binding of the carboxyl group to the amine creates an unstable complex, which results in cyclizing of the α-amino acid groups into an N-carboyxanhydride releasing HS^-^. Both reaction steps are reversible and the progression towards HS^-^ release only occurs in the presence of a sufficient bicarbonate concentration.

**Fig. 5.**
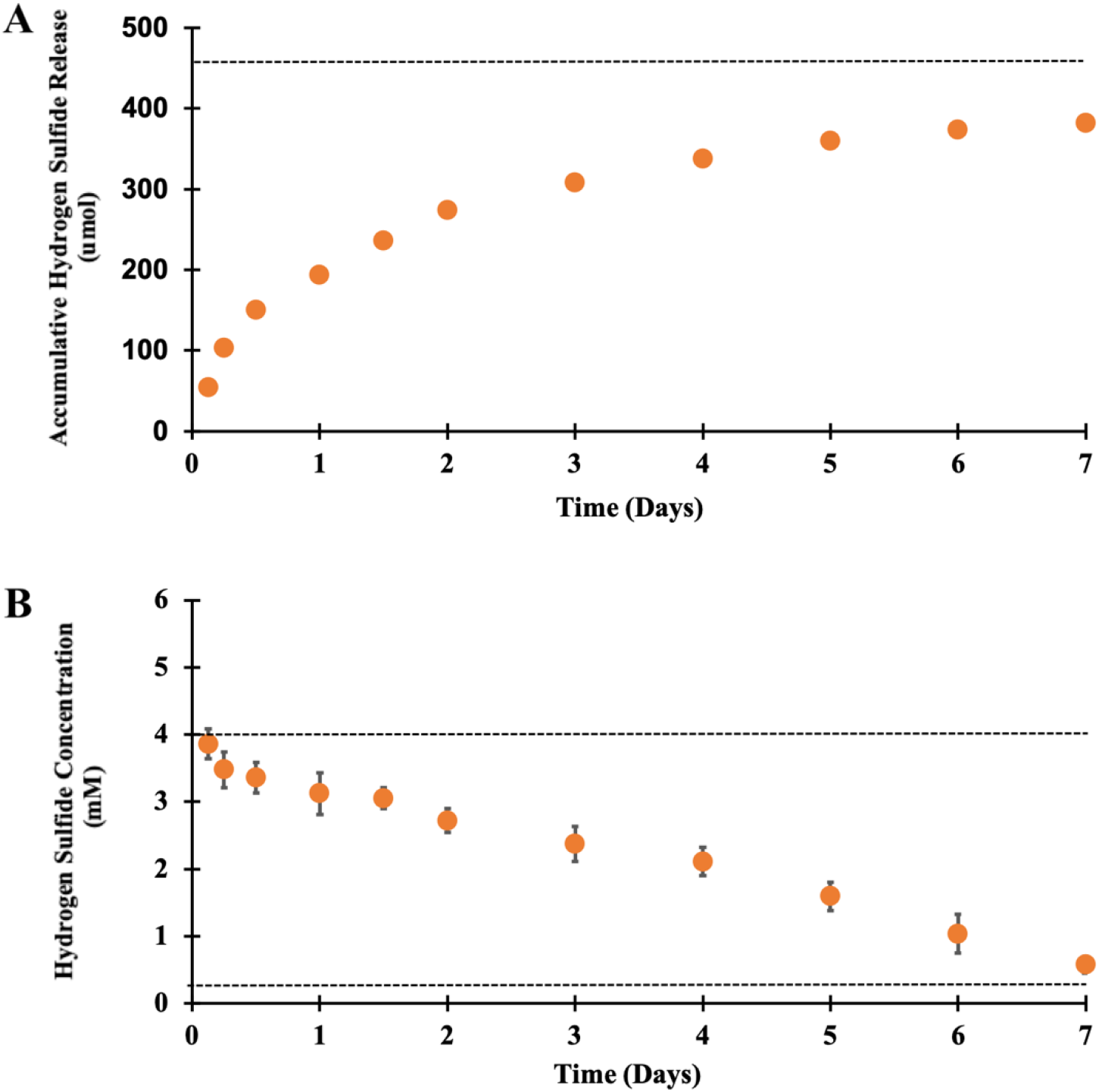
GluSH hydrogen sulfide release. **(a)** The cumulative release of H_2_S alleviated from 32 mM GluSH immersed in bicarbonate-supplemented Hank’s Balanced Salt Solution (HBSS) was measured over 7 days at 37 °C. **(b)** The solution H_2_S concentration was determined from this release data.

### GluSH Mediated Stem Cell Proliferation and Differentiation

To evaluate GluSH bioactivity, MSCs were exposed to GluSH_32_ supplemented with toxic level of Ca^2+^ and P_i_. To distinguish the difference between the effect of H_2_S release from the rest of GluSH molecule, glutamic acid (Glu) supplemented media was also investigated as a control. The proliferation and viability of cells exposed to media supplemented with excess Ca^2+^ and P_i_ with or without Glu or GluSH is summarized in **Fig. 6**. Proliferation of MSCs exposed to Ca^2+^ and/or P_i_ in the presence of GluSH (*i.e*., Ca_32_/GluSH_32_, P_16_/GluSH_32_, and Ca_32_/P_16_/GluSH_32_) were statistically significantly greater than comparative ones supplemented with Ca^2+^ and/or P_i_ alone or with Glu (*i.e*., Ca_32_, P_16_, Ca_32_/P_16_, Ca_32_/Glu_32_, P_16_/Glu_32_, and Ca_32_/P_16_/Glu_32_) and lower than media-only control and GluSH_32_ (**Fig. 6A** and **Table S5**). These results are due to the synergistic effect of H_2_S moderating high concentration Ca^2+^ and P_i_ cytotoxicity while allowing these molecules to likely carry out their osteoinductive effects. Interestingly, cells exposed to GluSH_32_ alone proliferated to 8 times of their initial seeding concentration indicating the biocompatibility of our novel molecule. However, MSCs exposed to Glu_32_ alone did not proliferate over the course of 14 days. This results suggest that the acidic nature of glutamic acid could potentially cause some cytotoxicity.^40^ In contrast, GluSH releasing HS^-^ produces the byproduct glutamate N-carboxyanhydride (**Fig. 5**), which generates a less acidic solution that glutamic acid while the presence of H_2_S would also serve as a cytoprotectant. The viability of the MSCs exposed to different combinations of signaling molecules reveals that the cells exposed to Ca_32_ and P_16_ as well as Glu were less than 30% metabolically active as those in the control group by day 14 (**Fig. 6B** and **Table S5**). Supplementing high Ca^2+^ and/or P_i_ concentration media with GluSH mitigated any cytotoxic effects increasing cell viability to more than 100% as compared to the control group.

**Fig. 6.**
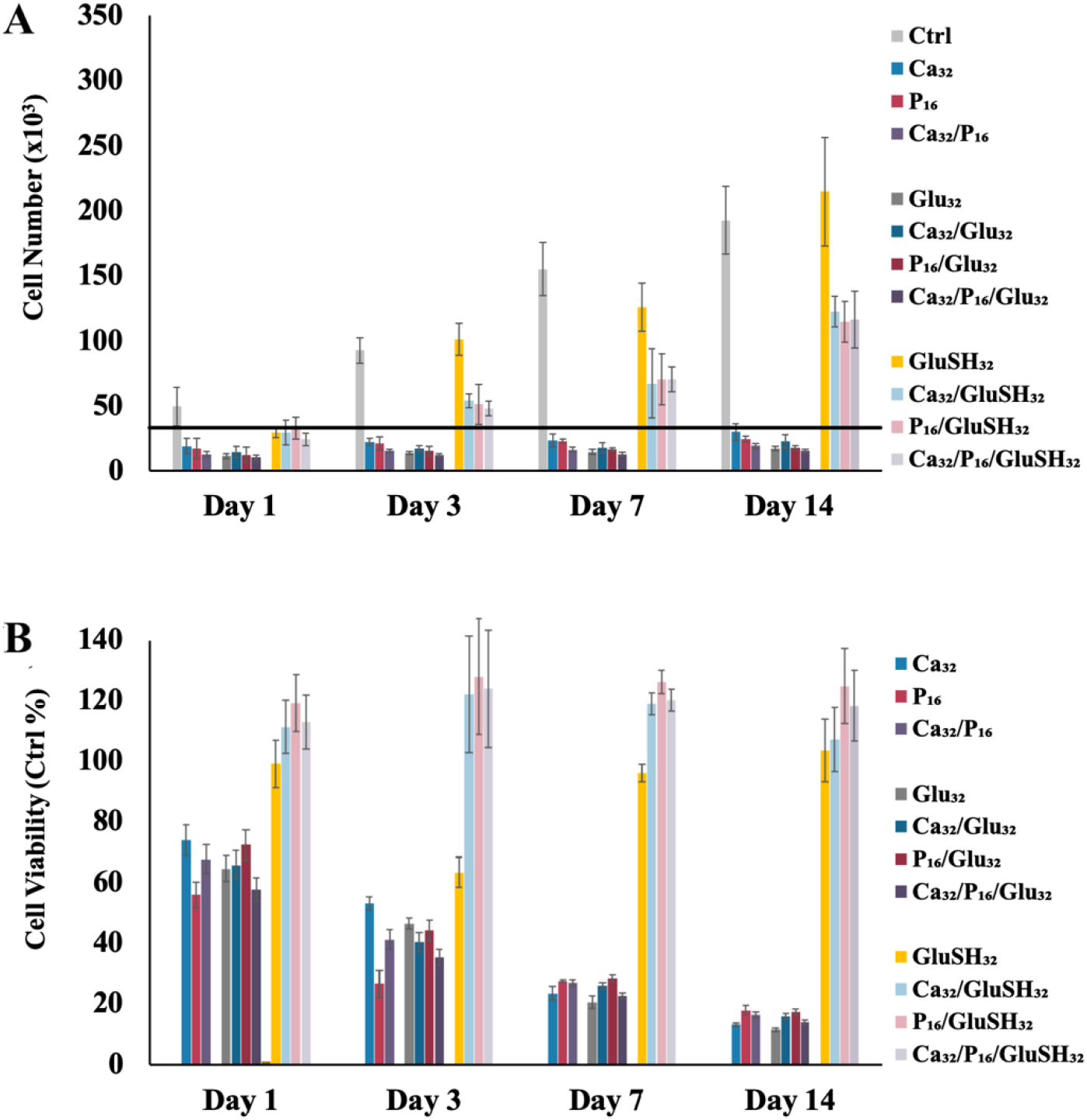
Proliferation and viability of MSCs exposed to different combinations of Ca^2+^, P_i_, and/or GluSH supplemented media. **(a)** Cell count was measured by the Quanti-iT PicoGreen Assay over 14 days of cell culture in media supplemented with no signaling molecules (Ctrl), 32 mM Ca^2+^ (Ca_32_), 16 mM P_i_ (P_16_), 32 mM glutamic acid (Glu_32_), 32 mM thioglutamic acid (GluSH_32_), or combination of these molecules (*i.e*., Ca_32_/P_16_, Ca_32_/Glu_32_, P_16_/Glu_32_, Ca_32_/P_16_/Glu_32_, Ca_32_/GluSH_32_, P_16_/GluSH_32_, Ca_32_/P_16_/GluSH_32_) for the first 7 days. This was followed by the cells being exposed for the next 7 days to no signaling molecules or half the Ca^2+^ and/or P_i_ concentrations (*i.e*., Ca_16_ and/or P_8_) they were originally cultured in. The black line indicates the original cell seeding number (*i.e*., 30,000). **(b)** An indirect measure of cell metabolism and viability was determined by NAD(P)H activity over 14 days using an MTS assay. MSCs were exposed to Ca_32_ and/or P_16_ with or without Glu_32_ or GluSH_32_ for the first 7 days followed by the aforementioned decrease in Ca^2+^ and/or P_i_ concentrations for the respective groups. All data were normalized against the results determined for MSCs given non-supplemented media. Statistical analysis of the data is available in the supplementary information (**Table S5** and **Table S6**).

The ALP activity and cell-based mineralization of MSCs cultured with Ca_32_ and/or P_16_ with or without GluSH_32_ or Glu_32_ is described in **Fig. 7**. MSCs exposed to non-supplemented media (Ctrl) or media supplemented with Glu_32_ or GluSH_32_ showed background levels of ALP expression while cells subjected to Ca_32_, P_16_, Ca_32_/P_16_, Ca_32_/GluSH_32_, P_16_/GluSH_32_, and Ca_32_/P_16_/GluSH_32_ showed statistically significant increased levels of ALP expression compared to control (**Fig. 7A**, **Table S5**). Cells cultured with Ca_32_ and P_16_ with GluSH_32_ (*i.e*., Ca_32_/GluSH_32_, P_16_/GluSH_32_, and Ca_32_/P_16_/GluSH_32_) showed statistically significantly higher levels of ALP expression compared to those without GluSH_32_ (*i.e*., Ca_32_, P_16_, and Ca_32_/P_16_) that also increased over time (**Fig. 7A**, **Table S5**, and **Table S6**). Assessment of MSC mineralization revealed that Ca_32_/GluSH_32_, P_16_/GluSH_32_, and Ca_32_/P_16_/GluSH_32_ induced statistically significant mineral deposition over time (**Table S6**) that was greater than all other treatments from Day 3 on (**Fig. 7B** and **Table S5**). MSCs exposed to Ca_32_, P_16_, or Ca_32_/P_16_ showed limited changes over time (**Table S6**) which while possessing greater mineralization than Ctrl was statistically insignificant different than cells exposed to Glu_32_ (**Fig. 7B** and **Table S5**).

**Fig. 7.**
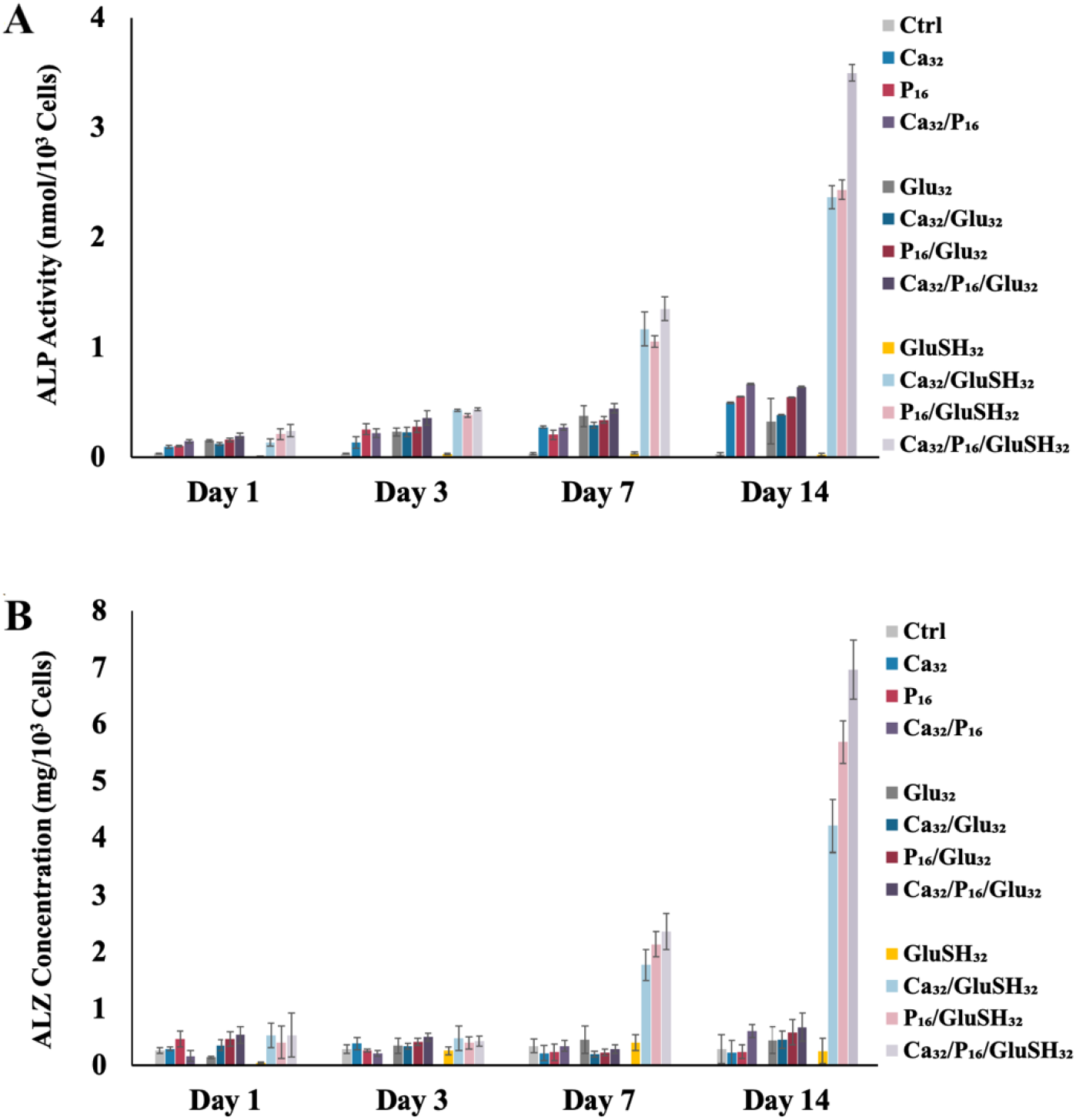
ALP activity and cell based mineralization of MSCs exposed to different combinations of Ca^2+^, P_i_, and/or GluSH supplemented media. **(a)** Maturation enzyme activity was analyzed by an ALP Assay Kit over 14 days of cell culture in media supplemented with no signaling molecules (Ctrl), 32 mM Ca^2+^ (Ca_32_), 16 mM P_i_ (P_16_), 32 mM glutamic acid (Glu_32_), 32 mM thioglutamic acid (GluSH_32_), or combination of these molecules (*i.e*., Ca_32_/P_16_, Ca_32_/Glu_32_, P_16_/Glu_32_, Ca_32_/P_16_/Glu_32_, Ca_32_/GluSH_32_, P_16_/GluSH_32_, and Ca_32_/P_16_/GluSH_32_) for the first 7 days. This was followed by the cells being exposed for the next 7 days to no signaling molecules or half the Ca^2+^ and/or P_i_ concentrations (*i.e*., Ca_16_ and/or P_8_) they were originally cultured in. **(b)** Alizarin red (ALZ) staining was used as an indirect measure of mineralization over 14 days of cell culture MSCs were exposed to Ca_32_ and/or P_16_ with or without Glu_32_ or GluSH_32_ for the first 7 days followed by the aforementioned decrease in Ca^2+^ and/or P_i_ concentrations for the respective groups. Both ALP activity and ALZ mineralization data were normalized on a per cell basis. Statistical analysis of the data is available in the supplementary information (**Table S5** and **Table S6**).

## CONCLUSIONS

This research aimed to evaluate the cytoprotective effect of H_2_S on MSCs experiencing ion overload similar to what can occur in a bone defect site microenvironment. It was determined that an effective therapeutic range exists for H_2_S that can improve cell proliferation and differentiation even in the presence of cytotoxic levels of calcium and phosphate ions. Furthermore, a novel compound was generated through glutamic acid modification that was capable of sustained H_2_S release in the therapeutic window for 7 days, which proved effective in mitigating ion-mediated cytotoxicity. These results support the considerable promise of thioglutamic acid as a useful product in helping to facilitate cell survival in critical sized bone defects.

## ACKNOWLEDGMENTS

The authors gratefully acknowledge support from start-up funds as well as a College of Engineering Incentive Fund Grant and a University of Missouri Research Council Grant, all kindly provided by the University of Missouri.

## REFERENCES

1. H. Wang, M. Naghavi, C. Allen, R. M. Barber, Z. A. Bhutta, A. Carter, D. C. Casey, F. J. Charlson, A. Z. Chen and M. M. Coates, The lancet, 2016, 388, 1459–1544.

2. S. Sheweita and K. Khoshhal, Current drug metabolism, 2007, 8, 519–525.

3. M. Marenzana and T.R. Arnett, Bone research, 2013, 1, 203.

4. K.A. Seta, Y. Yuan, Z. Spicer, G. Lu, J. Bedard, T.K. Ferguson, P. Pathrose, A. Cole-Strauss, A. Kaufhold and D.E. Millhorn, Cell Calcium, 2004, 36, 331–340.

5. Y. Sato, M. Kaji, F. Higuchi, I. Yanagida, K. Oishi and K. Oizumi, Osteoporosis international, 2001, 12, 445–449.

6. R. Marsell and T.A. Einhorn, Injury, 2011, 42, 551–555.

7. S. Orrenius and P. Nicotera, Journal of neural transmission. Supplementum, 1994, 43, 1–11.

8. J.-H. Zeng, S.-W. Liu, L. Xiong, P. Qiu, L.-H. Ding, S.-L. Xiong, J.-T. Li, X.-G. Liao and Z.-M. Tang, Journal of orthopaedic surgery and research, 2018, 13, 33.

9. R.Z. LeGeros, Clinical Orthopaedics and Related Research®, 2002, 395, 81–98.

10. M. Vallet-Regí and J. M. González-Calbet, Progress in Solid State Chemistry, 2004, 32, 1–31.

11. L. Wang and G. H. Nancollas, Chemical reviews, 2008, 108, 4628–4669.

12. J. Humeau, J. M. Bravo-San Pedro, I. Vitale, L. Nuñez, C. Villalobos, G. Kroemer and L. Senovilla, Cell Calcium, 2018, 70, 3–15.

13. Y. C. Chai, S. J. Roberts, E. Desmet, G. Kerckhofs, N. van Gastel, L. Geris, G. Carmeliet, J. Schrooten and F. P. Luyten, Biomaterials, 2012, 33, 3127–3142.

14. S. A. A. Ghavimi, B. N. Allen, J. L. Stromsdorfer, J.S. Kramer, X. Li and B.D. Ulery, Biomedical Materials, 2018, 13, 055005.

15. S. A. A. Ghavimi, E.S. Lungren, T.J. Faulkner, M.A. Josselet, Y. Wu, Y. Sun, F.M. Pfeiffer, C.L. Goldstein, C. Wan and B.D. Ulery, International Journal of Biological Macromolecules, 2019, 130, 88–98.

16. S. A. A. Ghavimi, E.S. Lungren, J.L. Stromsdorfer, B.T. Darkow, J.A. Nguyen, Y. Sun, F.M. Pfeiffer, C.L. Goldstein, C. Wan and B.D. Ulery, The AAPS Journal, 2019, 21, 41.

17. M. M. Gadalla and S.H. Snyder, Journal of neurochemistry, 2010, 113, 14–26.

18. Y. Kimura, Y.-I. Goto and H. Kimura, Antioxidants & redox signaling, 2010, 12, 1–13.

19. Z.-Z. Xie, Y. Liu and J.-S. Bian, Oxidative medicine and cellular longevity, 2016, 2016.

20. J. Yi, Y. Yuan, J. Zheng and T. Zhao, Journal of biochemical and molecular toxicology, 2019, 33, e22255.

21. B. Wu, H. Teng, L. Zhang, H. Li, J. Li, L. Wang and H. Li, Oxidative Medicine and Cellular Longevity, 2015, 2015.

22. G. Caliendo, G. Cirino, V. Santagada and J.L. Wallace, Journal of medicinal chemistry, 2010, 53, 6275–6286.

23. N. Shibuya, S. Koike, M. Tanaka, M. Ishigami-Yuasa, Y. Kimura, Y. Ogasawara, K. Fukui, N. Nagahara and H. Kimura, Nature communications, 2013, 4, 1366.

24. L. Li, M. Whiteman, Y.Y. Guan, K.L. Neo, Y. Cheng, S.W. Lee, Y. Zhao, R. Baskar, C.-H. Tan and P.K. Moore, Circulation, 2008, 117, 2351–2360.

25. W.-b. Wei, X. Hu, X.-d. Zhuang, L.-z. Liao and W.-d. Li, Molecular and cellular biochemistry, 2014, 389, 249–256.

26. Z. Zhou, M. von Wantoch Rekowski, C. Coletta, C. Szabo, M. Bucci, G. Cirino, S. Topouzis, A. Papapetropoulos and A. Giannis, Bioorganic & medicinal chemistry, 2012, 20, 2675–2678.

27. M. Bucci, A. Papapetropoulos, V. Vellecco, Z. Zhou, A. Pyriochou, C. Roussos, F. Roviezzo, V. Brancaleone and G. Cirino, Arteriosclerosis, thrombosis, and vascular biology, 2010, 30, 1998–2004.

28. A. Oshita, G. Rothstein and G. Lonngi, Blood, 1977, 49, 585–591.

29. A.A. Caro, S. Thompson and J. Tackett, Cell biology and toxicology, 2011, 27, 439.

30. M.S. Attene-Ramos, E.D. Wagner, H. R. Gaskins and M.J. Plewa, Molecular cancer research, 2007, 5, 455–459.

31. K.E. Benders, P. R. van Weeren, S.F. Badylak, D.B. Saris, W. J. Dhert and J. Malda, Trends in biotechnology, 2013, 31, 169–176.

32. Y. Luo, X. Liu, Q. Zheng, X. Wan, S. Ouyang, Y. Yin, X. Sui, J. Liu and X. Yang, Biochemical and biophysical research communications, 2012, 425, 473–477.

33. P. Varanyuwatana and A.P. Halestrap, Mitochondrion, 2012, 12, 120–125.

34. S. M. Schuster and M.S. Olson, Journal of Biological Chemistry, 1974, 249, 7159–7165.

35. J.W. Calvert, W. A. Coetzee and D.J. Lefer, Antioxidants & redox signaling, 2010, 12, 1203–1217.

36. V.A. Aleshin, A.V. Artiukhov, H. Oppermann, A.V. Kazantsev, N. V. Lukashev and V.I. Bunik, Cells, 2015, 4, 427–451.

37. F. Zhan, Y. Watanabe, A. Shimoda, E. Hamada, Y. Kobayashi and M. Maekawa, Clinica Chimica Acta, 2016.

38. L. Gambari, E. Amore, R. Raggio, W. Bonani, M. Barone, G. Lisignoli, B. Grigolo, A. Motta and F. Grassi, Materials Science and Engineering: C, 2019, 102, 471–482.

39. W. Chen, M. L. Melamed and M.K. Abramovitz, American Journal of Kidney Diseases, 2015, 65, 240–248.

40. S. A. A. Ghavimi, R.R. Tata, A.J. Greenwald, B.N. Allen, D.A. Grant, S.A. Grant, M. W. Lee and B.D. Ulery, The AAPS Journal, 2017, 19, 1029–1044.

