## Supplementary Information for "Controlled Hydrogen Sulfide Delivery to Enhance Cell Survival in Bone Tissue Engineering"

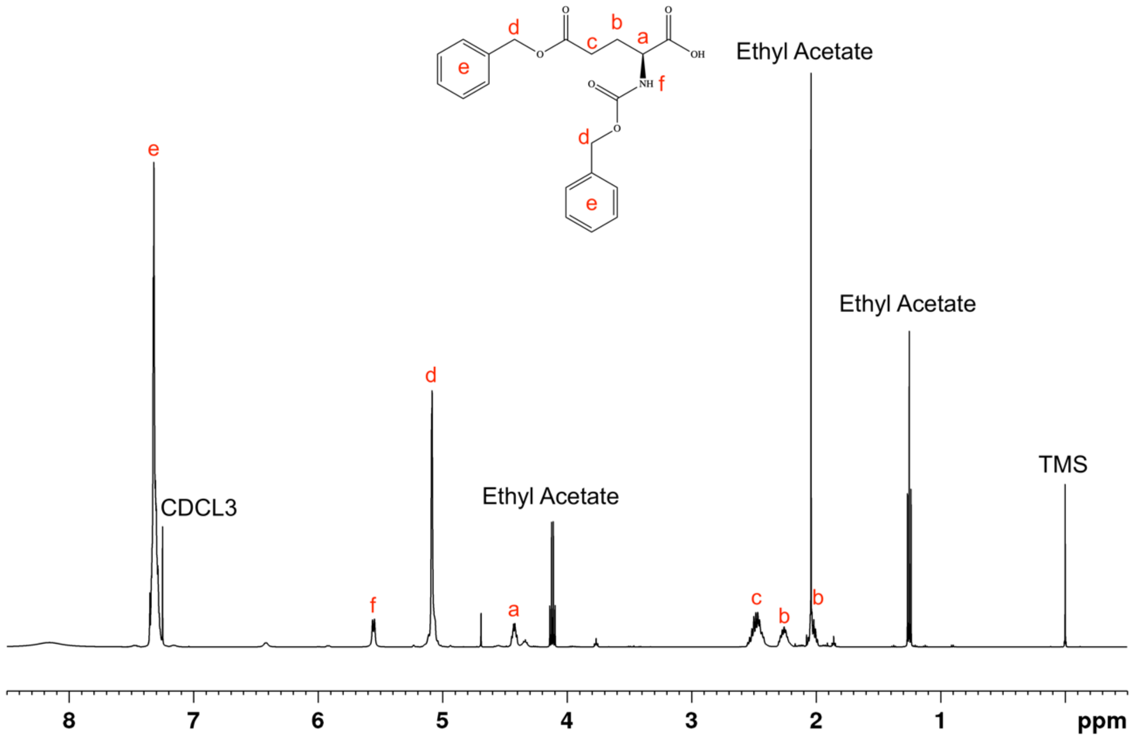


**Figure S1.** NMR spectrum of N-benzyloxycarbonyl-L-glutamic acid 5-benzyl ester.

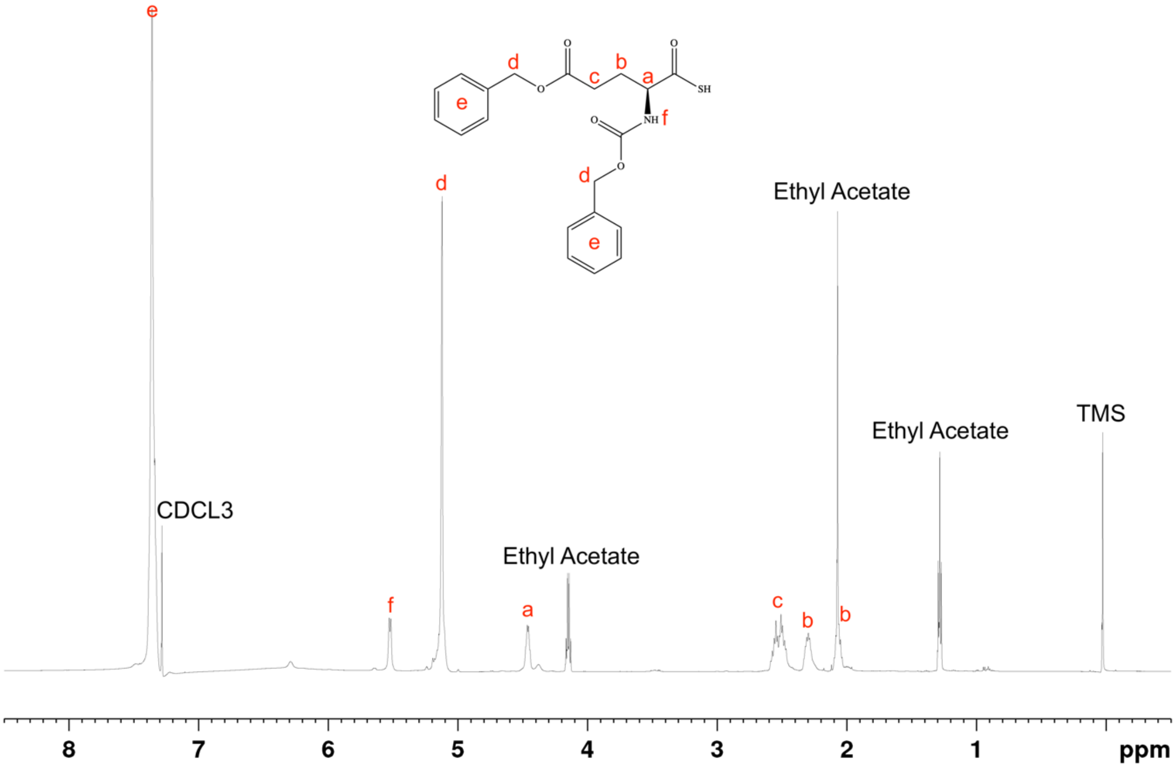


**Figure S2.** NMR spectrum of N-benzyloxycarbonyl-L-thioglutamic acid 5-benzyl ester.

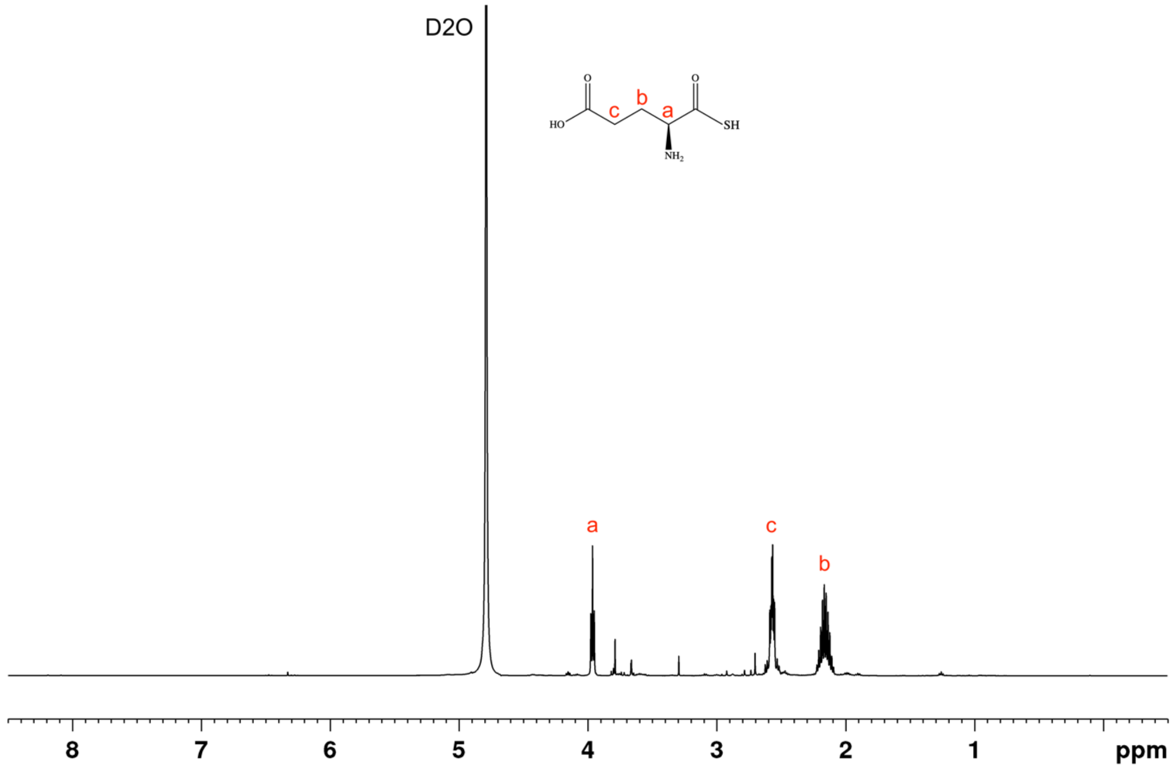


**Figure S3.** NMR spectrum of L-thioglutamic acid.

**Table S1.** Statistical analysis using Tukey's HSD test for data presented in **Figure 2** between different samples at the same time points. Groups that possess different letters have statistically significant differences ($p < 0.05$) in mean whereas those that possess the same letter are statistically similar.

| NaSH | Cell Number | MTS |
|---|---|---|
| Day 1 | | |
| Ctrl | A | |
| 0.25mM | AB | BCD |
| 0.5mM | ABC | ABCD |
| 1mM | ABC | D |
| 2mM | ABC | CD |
| 4mM | ABC | ABCD |
| 8mM | CD | A |
| 16mM | CD | ABC |
| 32mM | BCD | AB |
| 64mM | CD | AB |
| 128mM | D | AB |
| 256mM | D | AB |

| | | |
|---|---|---|
| Day 3 | | |
| Ctrl | AB | |
| 0.25mM | A | A |
| 0.5mM | A | A |
| 1mM | A | A |
| 2mM | A | A |
| 4mM | A | A |
| 8mM | BC | A |
| 16mM | C | A |
| 32mM | C | A |
| 64mM | C | A |
| 128mM | C | A |
| 256mM | C | A |
| Day 7 | | |
| Ctrl | AB | |
| 0.25mM | A | A |
| 0.5mM | A | A |
| 1mM | A | A |
| 2mM | AB | A |
| 4mM | AB | A |
| 8mM | BC | A |
| 16mM | C | A |
| 32mM | C | A |
| 64mM | C | A |
| 128mM | C | A |
| 256mM | C | A |
| Day 14 | | |
| Ctrl | B | |
| 0.25mM | AB | ABC |
| 0.5mM | A | ABCD |
| 1mM | A | ABCD |
| 2mM | AB | ABCD |
| 4mM | AB | AB |
| 8mM | C | A |
| 16mM | CD | ABCD |
| 32mM | C | CD |
| 64mM | DE | BCD |
| 128mM | E | BCD |
| 256mM | E | D |

**Table S2.** Statistical analysis using Tukey's HSD test for data presented in **Figure 2** and between the same samples at different time points. Groups that possess different letters have statistically significant differences ($p < 0.05$) in mean whereas those that possess the same letter are statistically similar.

| NaSH | Cell Number | MTS |
|---|---|---|
| Ctrl | | |
| Day 1 | Z | |
| Day 3 | Y | |
| Day 7 | X | |
| Day 14 | W | |
| 0.25mM | | |
| Day 1 | Z | Z |
| Day 3 | Y | Z |
| Day 7 | Y | Z |
| Day 14 | X | Z |
| 0.5mM | | |
| Day 1 | Z | Z |
| Day 3 | Y | Z |
| Day 7 | Y | Z |
| Day 14 | X | Z |
| 1mM | | |
| Day 1 | Z | Z |
| Day 3 | YZ | Z |
| Day 7 | Y | Z |
| Day 14 | X | Z |
| 2mM | | |
| Day 1 | Z | Z |
| Day 3 | Y | Z |
| Day 7 | Y | Z |
| Day 14 | X | Z |
| 4mM | | |
| Day 1 | Z | Z |
| Day 3 | Y | Z |
| Day 7 | Y | Z |
| Day 14 | X | Z |
| 8mM | | |
| Day 1 | Z | Z |
| Day 3 | Y | Z |
| Day 7 | Y | Z |
| Day 14 | X | Z |
| 16mM | | |
| Day 3 | Z | Z |
| Day 7 | Z | Z |
| Day 14 | Z | Z |
| Day 21 | Y | Z |

| | | |
|---|---|---|
| 32mM | | |
| Day 1 | Z | Z |
| Day 3 | Z | YZ |
| Day 7 | Z | Y |
| Day 14 | Y | Y |
| 64mM | | |
| Day 3 | Z | Y |
| Day 7 | YZ | YZ |
| Day 14 | XY | Y |
| Day 21 | X | Y |
| 128mM | | |
| Day 1 | Z | Z |
| Day 3 | Z | Y |
| Day 7 | Z | Z |
| Day 14 | Z | Y |
| 256mM | | |
| Day 1 | Z | Z |
| Day 3 | Z | Z |
| Day 7 | Z | Z |
| Day 14 | Z | Z |

**Table S3.** Statistical analysis using Tukey's HSD test for data presented in **Figure 2** and **Figure 3** between different samples at the same time points. Groups that possess different letters have statistically significant differences ($p < 0.05$) in mean whereas those that possess the same letter are statistically similar.

| | Cell Number | MTS | ALP | ALZ |
|---|---|---|---|---|
| Day 1 | | | | |
| Ctrl | A | | C | A |
| $Ca_{32}$ | C | C | B | A |
| $P_{16}$ | C | D | B | A |
| $Ca_{32}/P_{16}$ | C | CD | AB | A |
| $NaSH_1$ | AB | B | B | A |
| $Ca_{32}/NaSH_1$ | BC | AB | AB | A |
| $P_{16}/NaSH_1$ | BC | AB | A | A |
| $Ca_{32}/P_{16}/NaSH_1$ | BC | A | A | A |
| Day 3 | | | | |
| Ctrl | B | | D | AB |
| $Ca_{32}$ | D | B | C | A |
| $P_{16}$ | D | B | C | AB |
| $Ca_{32}/P_{16}$ | D | B | C | B |
| $NaSH_1$ | A | A | D | B |
| $Ca_{32}/NaSH_1$ | C | A | AB | B |
| $P_{16}/NaSH_1$ | C | A | B | AB |
| $Ca_{32}/P_{16}/NaSH_1$ | C | A | A | B |
| Day 7 | | | | |
| Ctrl | A | | D | C |
| $Ca_{32}$ | BC | D | C | C |
| $P_{16}$ | BC | D | C | C |
| $Ca_{32}/P_{16}$ | C | D | C | C |
| $NaSH_1$ | A | C | D | C |
| $Ca_{32}/NaSH_1$ | B | B | AB | AB |
| $P_{16}/NaSH_1$ | B | B | B | B |
| $Ca_{32}/P_{16}/NaSH_1$ | BC | A | A | A |
| Day 14 | | | | |
| Ctrl | B | | F | D |
| $Ca_{32}$ | D | B | E | D |
| $P_{16}$ | D | B | E | D |
| $Ca_{32}/P_{16}$ | D | B | D | D |
| $NaSH_1$ | A | A | F | D |
| $Ca_{32}/NaSH_1$ | C | A | C | C |
| $P_{16}/NaSH_1$ | C | A | B | B |
| $Ca_{32}/P_{16}/NaSH_1$ | C | A | A | A |

**Table S4.** Statistical analysis using Tukey's HSD test for data presented in **Figure 2** and **Figure 3** between the same samples at different time points. Groups that possess different letters have statistically significant differences ($p < 0.05$) in mean whereas those that possess the same letter are statistically similar.

| | Cell Number | MTS | ALP | ALZ |
|---|---|---|---|---|
| Ctrl | | | | |
| Day 1 | Z | | Z | Z |
| Day 3 | Y | | Z | Z |
| Day 7 | X | | Z | Z |
| Day 14 | X | | Z | Z |
| $Ca_{32}$ | | | | |
| Day 1 | Z | Z | Z | Z |
| Day 3 | Z | Y | Z | Z |
| Day 7 | Z | X | Y | Z |
| Day 14 | Z | W | X | Z |
| $P_{16}$ | | | | |
| Day 1 | Z | Z | Z | Z |
| Day 3 | Z | Y | Y | Z |
| Day 7 | Z | Y | Y | Z |
| Day 14 | Z | X | X | Z |
| $Ca_{32}/P_{16}$ | | | | |
| Day 1 | Z | Z | Z | Z |
| Day 3 | YZ | Y | Y | Z |
| Day 7 | YZ | X | X | Z |
| Day 14 | Y | W | W | Y |
| $NaSH_1$ | | | | |
| Day 1 | Z | Z | Z | Z |
| Day 3 | Y | Z | Y | Z |
| Day 7 | Y | Z | XY | Z |
| Day 14 | X | Z | X | Z |
| $Ca_{32}/NaSH_1$ | | | | |
| Day 1 | Z | Z | Z | Z |
| Day 3 | YZ | Z | Z | Z |
| Day 7 | Y | Z | Y | Y |
| Day 14 | X | Z | X | X |
| $P_{16}/NaSH_1$ | | | | |
| Day 1 | Z | Z | Z | Z |
| Day 3 | YZ | Z | Z | Z |
| Day 7 | Y | Z | Y | Y |
| Day 14 | X | Z | X | X |
| $Ca_{32}/P_{16}/NaSH_1$ | | | | |
| Day 1 | Z | Z | Z | Z |
| Day 3 | YZ | Z | Z | Z |
| Day 7 | Y | Z | Y | Y |
| Day 14 | X | Z | X | X |

**Table S5.** Statistical analysis using Tukey's HSD test for data presented in **Figure 5** and **Figure 6** between different samples at the same time points. Groups that possess different letters have statistically significant differences ($p < 0.05$) in mean whereas those that possess the same letter are statistically similar.

| | Cell Number | MTS | ALP | ALZ |
|---|---|---|---|---|
| Day 1 | | | | |
| Ctrl | A | | FG | AB |
| $Ca_{32}$ | BCDE | C | EF | AB |
| $P_{16}$ | BCDE | E | DE | AB |
| $Ca_{32}/P_{16}$ | CDE | CDE | CDE | AB |
| $Glu_{32}$ | E | CDE | BCDE | AB |
| $Ca_{32}/Glu_{32}$ | CDE | CDE | DE | AB |
| $P_{16}/Glu_{32}$ | DE | CD | BCD | AB |
| $Ca_{32}/P_{16}/Glu_{32}$ | E | DE | ABC | A |
| $GluSH_{32}$ | BCD | B | G | B |
| $Ca_{32}/GluSH_{32}$ | BC | AB | CDE | A |
| $P_{16}/GluSH_{32}$ | AB | A | AB | AB |
| $Ca_{32}/P_{16}/GluSH_{32}$ | BCDE | AB | A | A |
| Day 3 | | | | |
| Ctrl | A | | E | AB |
| $Ca_{32}$ | C | BC | D | AB |
| $P_{16}$ | C | D | C | AB |
| $Ca_{32}/P_{16}$ | C | BCD | CD | B |
| $Glu_{32}$ | C | BCD | C | AB |
| $Ca_{32}/Glu_{32}$ | C | BCD | C | AB |
| $P_{16}/Glu_{32}$ | C | BCD | BC | AB |
| $Ca_{32}/P_{16}/Glu_{32}$ | C | CD | AB | A |
| $GluSH_{32}$ | A | B | E | AB |
| $Ca_{32}/GluSH_{32}$ | B | A | A | AB |
| $P_{16}/GluSH_{32}$ | B | A | A | AB |
| $Ca_{32}/P_{16}/GluSH_{32}$ | B | A | A | AB |
| Day 7 | | | | |
| Ctrl | A | | F | C |
| $Ca_{32}$ | C | DE | DE | C |
| $P_{16}$ | C | D | E | C |
| $Ca_{32}/P_{16}$ | C | D | DE | C |
| $Glu_{32}$ | C | E | CD | C |
| $Ca_{32}/Glu_{32}$ | C | DE | CDE | C |
| $P_{16}/Glu_{32}$ | C | D | CDE | C |
| $Ca_{32}/P_{16}/Glu_{32}$ | C | DE | C | C |
| $GluSH_{32}$ | A | C | F | C |
| $Ca_{32}/GluSH_{32}$ | B | B | B | B |
| $P_{16}/GluSH_{32}$ | B | A | B | AB |
| $Ca_{32}/P_{16}/GluSH_{32}$ | B | B | A | A |

| | | | | |
|---|---|---|---|---|
| Day 14 | | | | |
| Ctrl | A | | F | D |
| $Ca_{32}$ | C | C | CDE | D |
| $P_{16}$ | C | C | CD | D |
| $Ca_{32}/P_{16}$ | C | C | C | D |
| $Glu_{32}$ | C | C | E | D |
| $Ca_{32}/Glu_{32}$ | C | C | DE | D |
| $P_{16}/Glu_{32}$ | C | C | CD | D |
| $Ca_{32}/P_{16}/Glu_{32}$ | C | C | C | D |
| $GluSH_{32}$ | A | B | F | D |
| $Ca_{32}/GluSH_{32}$ | B | B | B | C |
| $P_{16}/GluSH_{32}$ | B | A | B | B |
| $Ca_{32}/P_{16}/ GluSH_{32}$ | B | AB | A | A |

**Table S6.** Statistical analysis using Tukey's HSD test for data presented in **Figure 5** and **Figure 6** between the same samples at different time points. Groups that possess different letters have statistically significant differences ($p < 0.05$) in mean whereas those that possess the same letter are statistically similar.

| | Cell Number | MTS | ALP | ALZ |
|---|---|---|---|---|
| Ctrl | | | | |
| Day 1 | Z | | Z | Z |
| Day 3 | Y | | Z | Z |
| Day 7 | X | | Z | Z |
| Day 14 | X | | Z | Z |
| $Ca_{32}$ | | | | |
| Day 1 | Z | Z | Z | Z |
| Day 3 | Z | Y | Z | Z |
| Day 7 | Z | X | Y | Z |
| Day 14 | Z | W | X | Z |
| $P_{16}$ | | | | |
| Day 1 | Z | Z | Z | Z |
| Day 3 | Z | Y | Y | Z |
| Day 7 | Z | Y | Y | Z |
| Day 14 | Z | X | X | Z |
| $Ca_{32}/P_{16}$ | | | | |
| Day 1 | Z | Z | Z | Z |
| Day 3 | YZ | Y | Y | Z |
| Day 7 | YZ | X | X | Z |
| Day 14 | Y | W | W | Y |
| $Glu_{32}$ | | | | |
| Day 1 | Z | Z | Z | Z |
| Day 3 | YZ | Y | Z | Z |
| Day 7 | YZ | X | Z | Z |
| Day 14 | Y | W | Z | Z |
| $Ca_{32}/Glu_{32}$ | | | | |
| Day 1 | Z | Z | Z | Z |
| Day 3 | Z | Y | Y | YZ |
| Day 7 | Z | X | X | YZ |
| Day 14 | Z | W | W | Y |
| $P_{16}/Glu_{32}$ | | | | |
| Day 1 | Z | Z | Z | Z |
| Day 3 | Z | Y | Y | YZ |
| Day 7 | Z | X | Y | YZ |
| Day 14 | Z | W | X | Y |
| $Ca_{32}/P_{16}/Glu_{32}$ | | | | |
| Day 1 | Z | Z | Z | Z |
| Day 3 | YZ | Y | Y | YZ |
| Day 7 | YZ | X | X | YZ |
| Day 14 | Y | W | W | Y |

| | | | | |
|---|---|---|---|---|
| $GluSH_{32}$ | | | | |
| Day 1 | Z | Z | Z | Z |
| Day 3 | Y | Z | YZ | YZ |
| Day 7 | Y | Z | Y | YZ |
| Day 14 | X | Z | Y | Y |
| $Ca_{32}/GluSH_{32}$ | | | | |
| Day 1 | Z | Z | Z | Z |
| Day 3 | YZ | Z | Y | Z |
| Day 7 | Y | Z | X | Y |
| Day 14 | X | Z | W | W |
| $P_{16}/GluSH_{32}$ | | | | |
| Day 1 | Z | Z | Z | Z |
| Day 3 | YZ | Z | Y | Z |
| Day 7 | Y | Z | X | Y |
| Day 14 | X | Z | W | W |
| $Ca_{32}/P_{16}/GluSH_{32}$ | | | | |
| Day 1 | Z | Z | Z | Z |
| Day 3 | YZ | Z | Y | Z |
| Day 7 | Y | Z | X | Y |
| Day 14 | X | Z | W | W |
